# Optimized pseudotyping conditions for the SARS-COV2 Spike glycoprotein

**DOI:** 10.1101/2020.05.28.122671

**Authors:** Marc C. Johnson, Terri D. Lyddon, Reinier Suarez, Braxton Salcedo, Mary LePique, Maddie Graham, Clifton Ricana, Carolyn Robinson, Detlef G. Ritter

## Abstract

The SARS-COV2 Spike glycoprotein is solely responsible for binding to the host cell receptor and facilitating fusion between the viral and host membranes. The ability to generate viral particles pseudotyped with SARS-COV2 Spike is useful for many types of studies, such as characterization of neutralizing antibodies or development of fusion-inhibiting small molecules. Here we characterized the use of a codon-optimized SARS-COV2 Spike glycoprotein for the generation of pseudotyped HIV-1, MLV, and VSV particles. The full-length Spike protein functioned inefficiently with all three systems but was enhanced over 10-fold by deleting the last 19 amino acids of the cytoplasmic tail of Spike. Infection of 293FT target cells was only possible if the cells were engineered to stably express the human ACE-2 receptor, but stably introducing an additional copy of this receptor did not further enhance susceptibility. Stable introduction of the Spike activating protease TMPRSS2 further enhanced susceptibility to infection by 5-10 fold. Substitution of the signal peptide of the Spike protein with an optimal signal peptide did not enhance or reduce infectious particle production. However, modification of a single amino acid in the furin cleavage site of Spike (R682Q) enhanced infectious particle production another 10-fold. With all enhancing elements combined, the titer of pseudotyped particles reached almost 10^6^ infectious particles/ml. Finally, HIV-1 particles pseudotyped with SARS-COV2 Spike was successfully used to detect neutralizing antibodies in plasma from COVID-19 patients, but not plasma from uninfected individuals.

**Importance:** When working with pathogenic viruses, it is useful to have rapid quantitative tests for viral infectivity that can be performed without strict biocontainment restrictions. A common way of accomplishing this is to generate viral pseudoparticles that contain the surface glycoprotein from the pathogenic virus incorporated into a replication-defective viral particle that contains a sensitive reporter system. These pseudoparticles enter cells using the glycoprotein from the pathogenic virus leading to a readout for infection. Conditions that block entry of the pathogenic virus, such as neutralizing antibodies, will also block entry of the viral pseudoparticles. However, viral glycoproteins often are not readily suited for generating pseudoparticles. Here we describe a series of modifications that result in the production of relatively high titer SARS-COV2 pseudoparticles that are suitable for detection of neutralizing antibodies from COVID-19 patients.

## Introduction

The COVID-19 pandemic is a severe threat to human health and the global economy. COVID-19 is caused by infection with the SARS-CoV-2 virus, which is a highly pathogenic betacoronavirus (1, 2). A critical tool for the study of pathogenic viruses such as SARS-COV2 is a rapid and sensitive assay for agents that block viral entry such as neutralizing antibodies or small molecule inhibitors. A safe and powerful technique for generating such an assay is to generate viral pseudotyped particles where the surface fusion protein of the pathogen of interest is assembled onto the surface of replication-defective virus which contains a sensitive reporter protein.

The Spike glycoprotein from Coronaviruses facilitates binding to the host cell receptor and fusion between viral and cellular membranes. The SARS-COV2 Spike protein is a large 1,274 amino acid protein that contains an N-terminal S1 receptor binding domain and a C-terminal S2 fusion domain. Fusion by SARS-COV2 Spike requires binding to the host receptor ACE2 (3-6) as well as proteolytic cleavage by host proteases at the S1/S2 and S2’ positions by host cysteine proteases cathepsin B and L (CatB/L) or serine protease TMPRSS2 (3, 6). Depending on the cell line, inhibition of one or both of these proteases is sufficient to block viral entry. An interesting difference between the SARS-COV1 and COV2 Spike proteins is the presence of a furin cleavage site near the S1/S2 cleavage site (6, 7). This cleavage site was found to be essential for infection of human lung cells (3). Surprisingly, passaging of SARS-COV2 in Vero E6 cells selects for mutations that alter the furin cleavage site, resulting in virus that produce larger plaque sizes (8).

There have been several reports generating SARS-COV1 (9-12) and SARS-COV2 (3-5, 13, 14) viral pseudotypes with glycoprotein-defective MLV, HIV, and VSV particles. Although pseudoparticles could be generated with full-length SARS-COV1 Spike, the pseudotyping efficiency was shown to be enhanced by about 100-fold by deleting the last 19 amino acids of the cytoplasmic tail (9) which removed a presumptive ER retention sequence. Mutation of the presumptive ER retention site in the cytoplasmic tail of SARS-COV2 (K1269A, H1271A) was not found to enhance pseudotyping efficiency (14). Here we develop optimized genetic conditions for generating a SARS-COV2 Spike pseudotyped particles.

## Results

### Truncation of SARS-COV2 Spike enhances viral pseudotyping

A codon optimized full length Spike gene was synthesized and introduced in place of the GFP gene in the plasmid pEGFP-N1. Because truncation of the cytoplasmic tail of SARS-COV1 Spike was shown to enhance production of viral pseudotypes (9), we also generated a SARS-COV2 Spike subclone with the last 19 amino acids deleted (Fig. 1). Pseudotyped particles were generated with the two Spike proteins as well as VSV-G with glycoprotein-defective MLV, HIV-1 and VSV particles (Fig. 1). To avoid potential false positive signal, a reporter system was utilized where the Cre recombinase gene is expressed by HIV-1 and MLV particles. When these particles transduce the Cre reporter cell line, the protein causes a recombination in an engineered reporter element that results in GFP expression. Because the viral producing cells do not express GFP, they are eliminated as a source of false positive signal. This 293FT based reporter cell line was further engineered to express human ACE2. Infectious particle production with all three types of particles was very low with the full-length Spike protein, but the Δ19 Spike produced viral titers approaching 10^4^ infectious particles/ml with both HIV-1 and VSV particles. The infectious particle production remained over 1000-fold lower than with the control glycoprotein, VSV-G.

**Figure 1.**
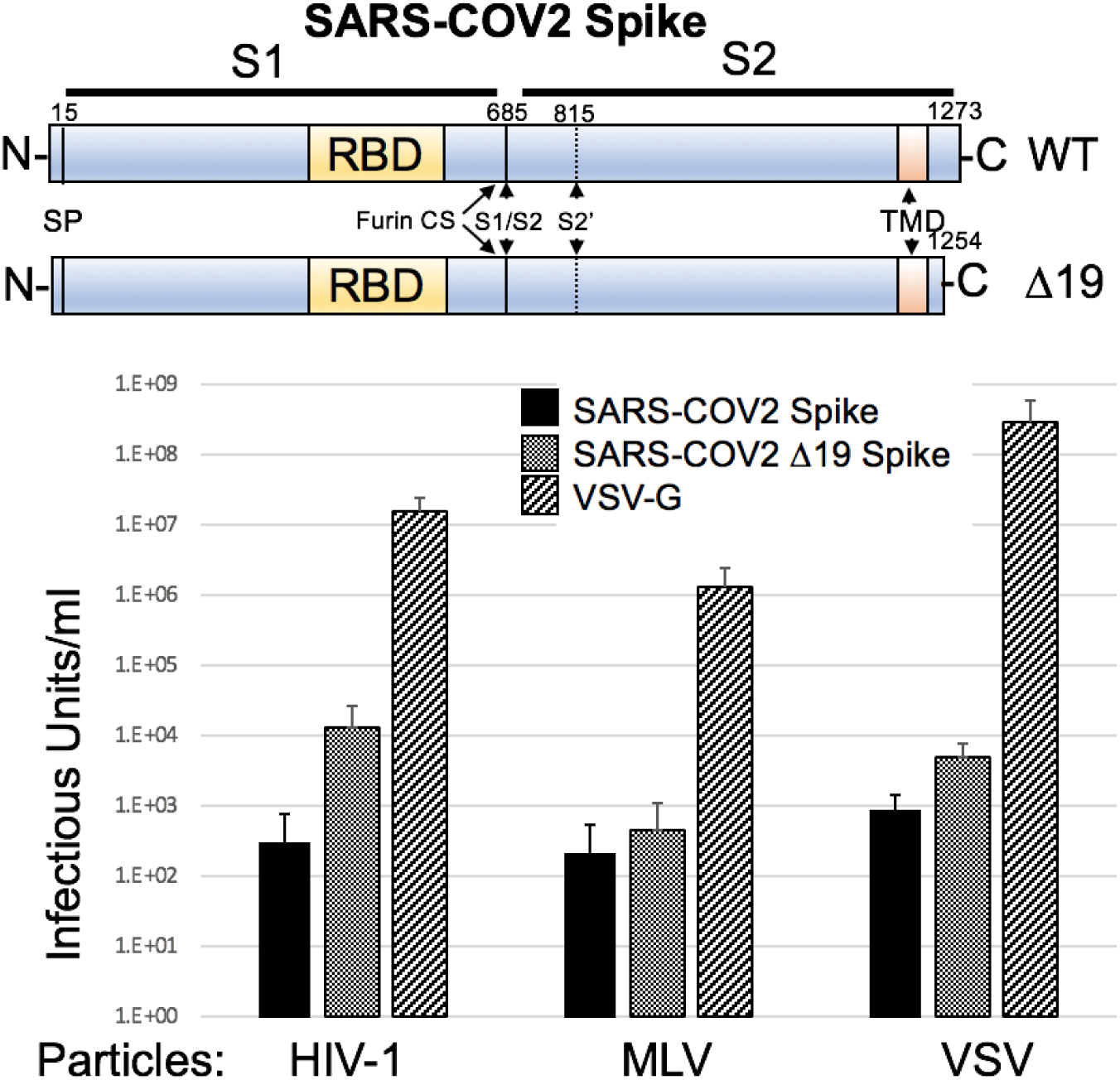
SARS-COV2 pseudotyped particles. Upper, schematic of SARS-COV2 Spike protein. RBD = receptor binding domain, SP = signal peptide, TMD = transmembrane domain, furin CS = furin cleavage site. Lower, infectious particle production of glycoprotein defective HIV-1, MLV, or VSV pseudoparticles with SARS-COV2 Spike, SARS-COV2 Δ19 Spike, and VSV-G. HIV-1 and MLV particles contained a Cre reporter and were scored on 293FT/ACE2 cells containing a Cre-inducible GFP reporter. VSV-G particles directly contained GFP (VSVΔG-GFP). Data are the average and standard deviation of three independent experiments.

To allow rapid quantitation with minimum background, we proceeded with an Env-defective HIV-1 provirus containing a *gausia* luciferase (Gluc) gene in the reverse orientation containing a forward intron (Fig. 2 (15)). This viral construct produces very low background because it is not capable of producing luciferase signal unless the gene is reverse transcribed in the target cell. Infectivity with SARS-COV2 Spike requires expression of the host receptor ACE2. Our starting cell line was generated using a retroviral transfer vector containing ACE2 and a puromycin selection cassette. To determine if introduction of an additional stable copy of ACE2 would enhance susceptibility to infection, we generated a second retroviral transfer vector with a blasticidin resistance cassette. Retroviral particles were generated with this vector and used to stably transduce 293FT or 293FT/ACE2 (puro) cells. Each of the cell lines was transduced with HIV-1-Gluc particles pseudotyped with SARS-COV2 Δ19 Spike (Fig. 2). As expected, 293FT cells were not susceptible to infection with the HIV-1/SARS-COV2 Δ19 Spike. However, both cell lines containing a single introduction of ACE2, and the cell line containing two introductions had approximately 1000-fold increase in luciferase signal, but the signal was roughly equivalent among the three cell lines.

**Figure 2.**
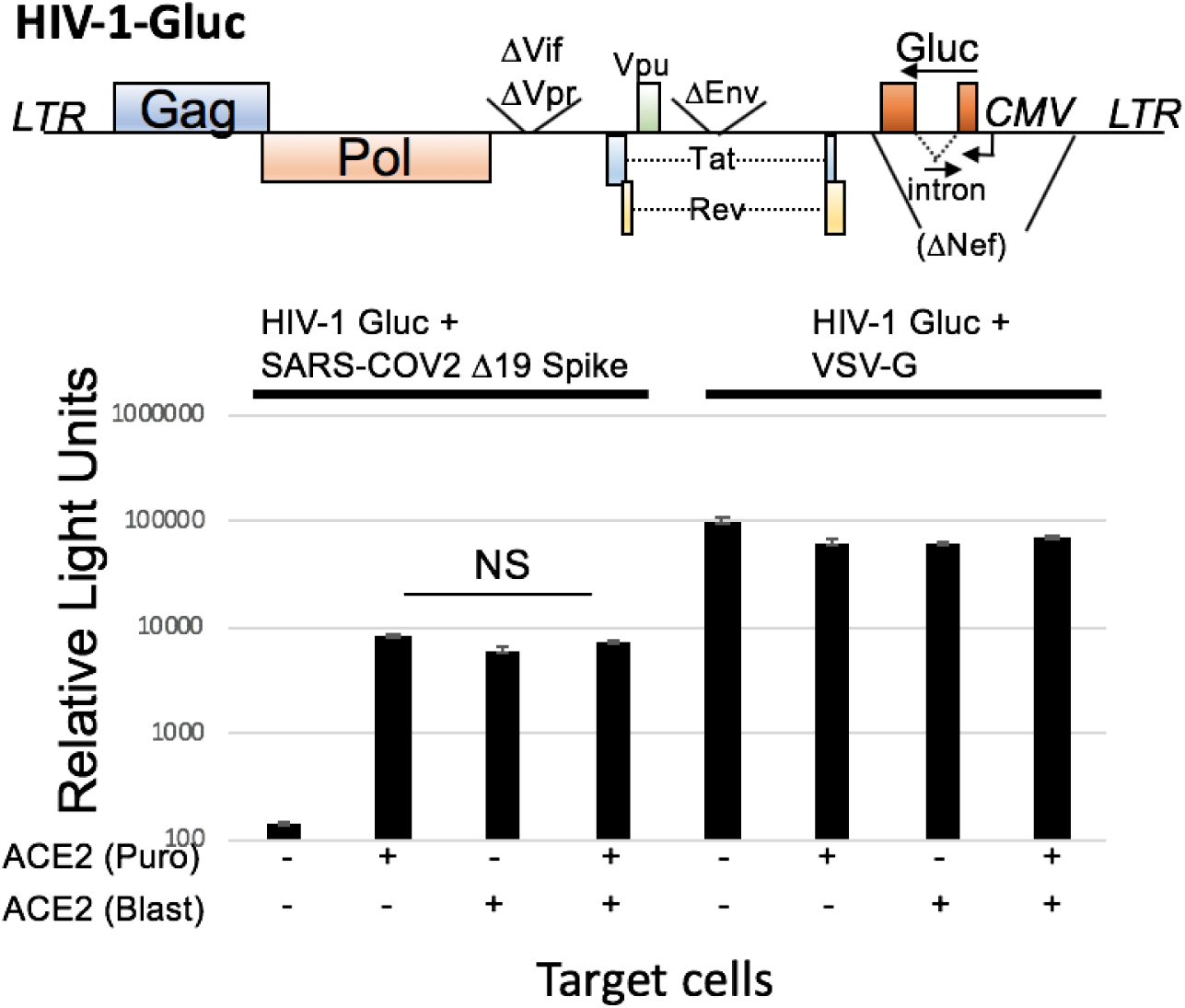
Introduction of ACE2 is required for infection of 293FT cells with SARS-COV2 Spike. Upper, schematic of the HIV-1 *gausia* luciferse vector. Lower, transduction of 293FT cells transduced with different numbers of ACE2 genes with HIV-1-Gluc pseudoparticles pseudotyped with SARS-COV2 Δ19 Spike or VSV-G. Transductions with VSV-G used 100-times less viral supernatant. Data are the average and standard deviation of a representative experiment performed in triplicate. NS = no significant difference.

### TMPRSS2 expression enhances susceptibility of cells to SARS-COV2 pseudotypes

The SARS-COV2 Spike requires proteolytic priming during infection by either cysteine proteases CatB/L or serine protease TMPRSS2 produced in the target cell (3, 6). To determine whether introduction of TMPRSS2 would enhance susceptibility to infection, we synthesized a codon-optimized TMPRSS2 gene, introduced this gene into a retroviral transfer vector, and stably transduced 293FT or 293FT/ACE2 cells. The stable introduction of TMPRSS2 to 293FT cells did not impart sensitivity to transduction with HIV-1 particles pseudotyped with SARS-COV2 Δ19 Spike (Fig. 3). However, stable introduction of TMPRSS2 to 293FT/ACE2 cells increased the Gluc signal from target cells by 5-10 fold. Neither ACE2 expression nor TMPRSS2 expression affected Gluc signal from particles pseudotyped with VSV-G.

**Figure 3.**
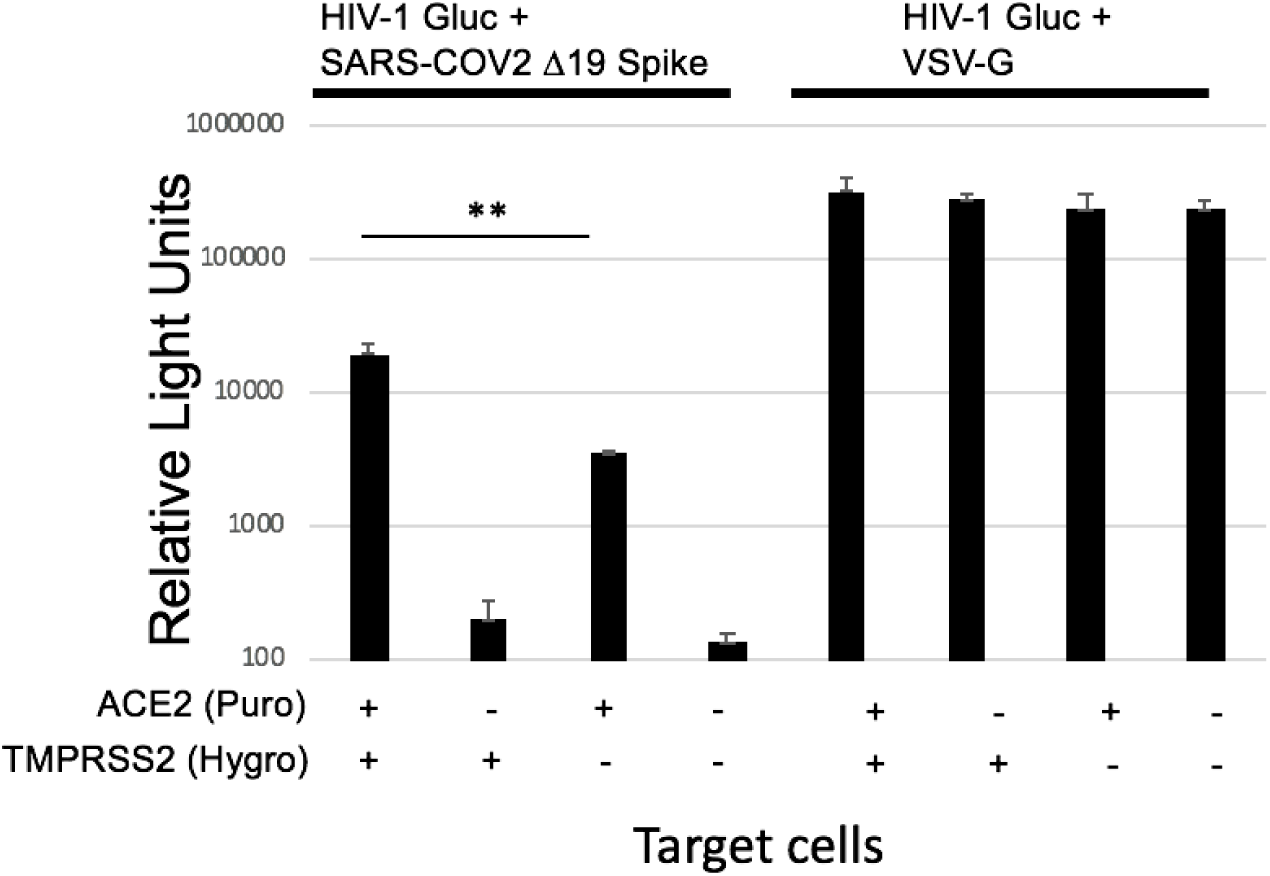
TMPRSS2 expression enhances susceptibility to infection with SARS-COV2 Spike. 293FT cells stably expressing ACE2, TMPRSS2, or both were transduced with HIV-1-Gluc particles pseudotyped with SARS-COV2 Δ19 Spike or VSV-G. Transductions with VSV-G used 100-times less viral supernatant. Data are the average and standard deviation of a representative experiment performed in triplicate. ** p<0.01 in paired Student’s t-test.

### The R682Q mutation in SARS-COV2 Spike enhances pseudotyping efficiency

To explore additional genetic modifications in SARS-COV2 that might enhance infectious particle production, we replaced the endogenous signal peptide with a well characterized strong signal peptide (16), and separately we introduced a point mutation in the furin-cleavage site of SARS-COV2 Δ19 Spike (R682Q) that has previously been reported to increase SARS-COV2 plaque sizes (Fig. 4 (8)). Infectious particle production was assessed using both Gluc and Cre reporters, and transductions were performed in cell lines with and without TMPRSS2. To minimize target cell variation, the transductions were all performed in 293FT/Cre-reporter/ACE2 and 293FT/Cre-reporter/ACE2/TMPRSS2 cells. Replacement of the signal peptide did not increase or decrease infectious particle production (Fig 4A). However, the R682Q mutation significantly enhanced infectious particle production under all conditions. The enhancement was most pronounced with virus containing the Cre reporter in cells lacking TMPRSS2 (11-fold). The highest titer of 8×10^5^ was achieved with the R682Q Δ19 Spike into cells expressing both ACE2 and TMPRSS2.

**Figure 4.**
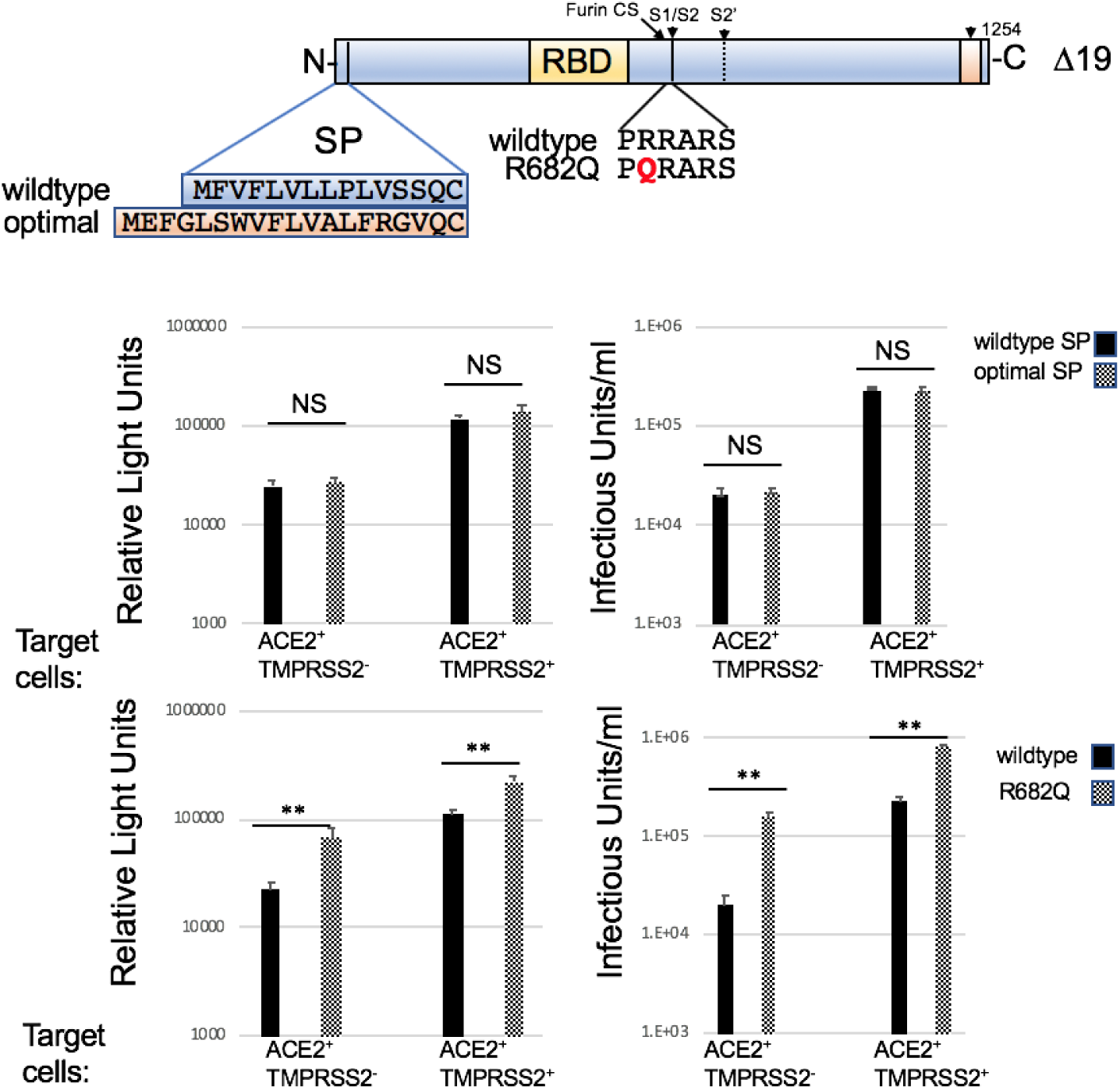
R682 mutation in SARS-COV2 Δ19 Spike enhances pseudotyping. Upper, schematic illustrating introduction of the optimal SP and R682Q into SARS-COV2 Δ19 Spike. HIV-1-GLuc and HIV-CMV-Cre were pseudotyped with SARS-COV2 Δ19 Spike, SARS-COV2 Δ19 Spike with optimal SP, and SARS-COV2 R682Q Δ19 Spike. Transductions were performed on 293FT/ACE2 and 293FT/ACE2/TMPRSS2 cells both containing a Cre-inducible GFP reporter. Data are the average and standard deviation of a representative experiment performed in triplicate. NS = no significant difference., ** p<0.01 in paired Student’s t-test.

**Figure 5.**
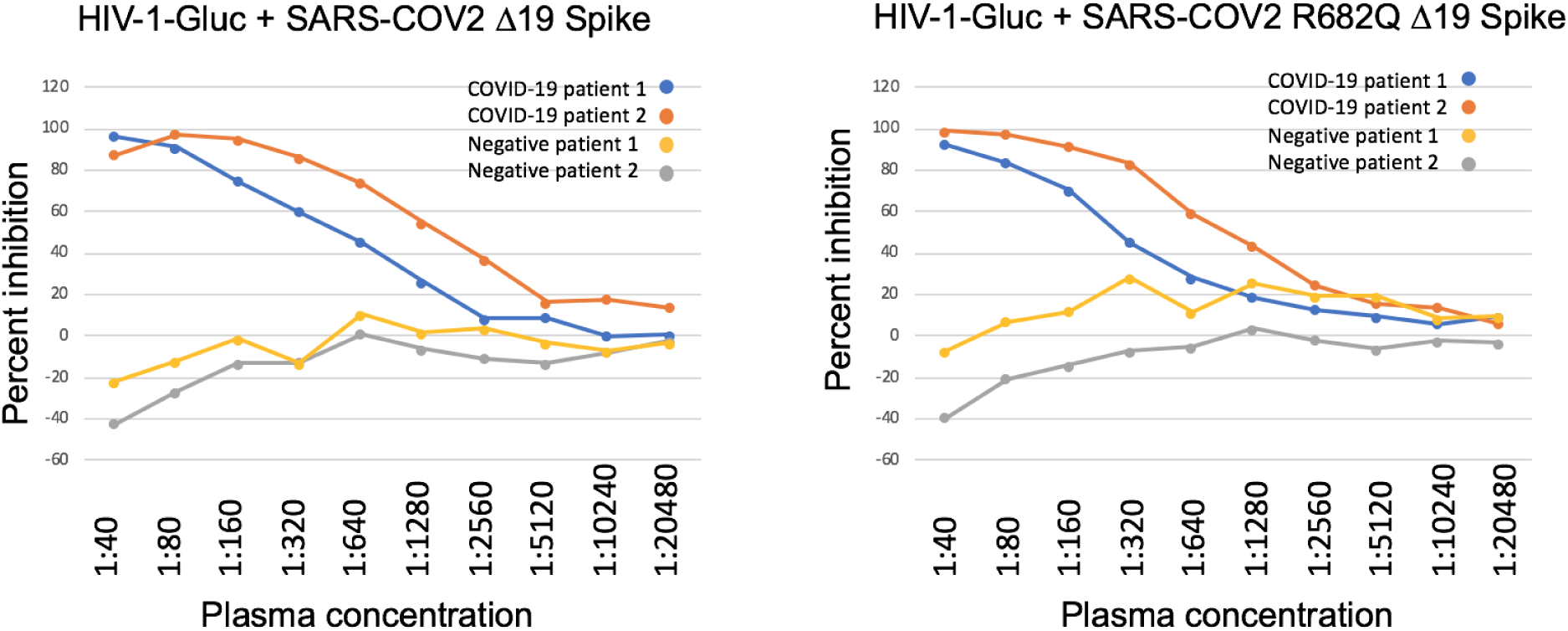
Particles pseudotyped with SARS-COV2 Spike function in neutralization assays. HIV-1-Gluc particles were pseudotyped with SARS-COV2 Δ19 Spike (left) or SARS-COV2 R682Q Δ19 Spike (right). Plasma was obtained from two patients with a confirmed COVID-19 infection, or infection negative patients. Two-fold serially diluted plasma was incubated with pseudotypes for 1 hour at 37C. The mixture was subsequently incubated with 293/ACE2/TMPRSS2 cells and Gluc measurements were taken 2 days post transduction.

### SARS-COV2 Spike pseudotypes function in neutralization assays

To test if the pseudoparticles produced were suitable for use in a neutralization assay, we incubated HIV-1 particles pseudotyped with SARS-COV2 Δ19 Spike with and without the R682Q mutation with serial dilutions of control plasma and plasma from COVID-19 patients before adding 293FT/ACE2/TMPRSS2 cells. Robust and quantifiable neutralization was detected in plasma from COVID-19 plasma, but not from control plasma. The R682Q mutation did not noticeably affect sensitivity to neutralizing antibodies.

## Methods

### Plasmids

The gammaretroviral transfer plasmids pQCXIP (puromycin resistance) and pQCXIH (pygromycin resistance) were obtained from Clontech. The blasticidin gammaretroviral transfer vector (pQCXIB) was generated by replacing the puromycin resistance cassette from PQCXIP with a blasticidin resistance cassette. The gammaretroviral vector for generating the Cre sensor cell line (MLV lox-mTomato-lox-GFP-Blast) was engineered into pQCXIB and contained the monomeric Tomato gene flanked by Lox sequences (ATAACTTCGTATAGCATACATTATACGAAGTTAT), which was followed immediately by eGFP. The ACE2 transfer vectors were generated by engineering the human ACE2 gene (NCBI Reference Sequence: NM_021804.3) into pQCXIP and pQCXIH. The TMPRSS2 transfer vector was generated by synthesizing a codon-optimized version of the human TMPRSS2 gene (NCBI Reference Sequence: NP_001128571.1) and engineering it into the vector pQCXIH. The HIV-1-CMV-Cre vector was an NL4-3 derived provirus that is defective in Vif, Vpr, Env and has the Nef gene replaced with CMV-Cre. The plasmid was generated by replacing the GFP gene from the previously described plasmid HIV-CMV-GFP+Vpu (17) with the Cre gene from plasmid MLV-Cre (18), provided by Alan Rein (NCI Frederick). The MLV+ CMV Cre vector was generated by replacing the GFP gene from MLV-GFP (provided by Shan-Lu Liu, The Ohio State University) with the Cre gene from MLV-Cre (18), provided by Alan Rein (NCI Frederick). The MLV GagPol expression construct was provided by Walther Mothes (Yale University). The HIV-1-Gluc vector was previously described (19). The VSV-G expression construct was obtained from the NIH AIDS Reagent Program (20). The SARS-COV2 expression vector was generated by synthesizing a codon-optimized version of the SARS-COV2 Spike (GenBank accession number MN985325.1). The Δ19 Spike and the Δ19 R682Q Spike were generated by PCR mutagenesis. The Spike subclone with the alternative signal peptide contained the previously described H7 peptide (16) was generated by synthesizing the 5’ end of the Spike gene with the alternative signal peptide and introducing it into the SARS-COV2 Δ19 Spike clone.

### Virus production and infectivity assays

All transfections were performed in 6 well plates. 293FT cells were transfected with a total of 1 microgram of plasmid and 4 micrograms of PEI (21). For HIV-1-CMV-Cre particles, cells were transfected with 900ng of provirus and 100ng of glycoprotein expression vector in Fig. 1, and 800ng of provirus and 200ng of glycoprotein expression vector in subsequent figures. For HIV-1-Gluc particles, cells were transfected with 800ng of provirus and 200ng of glycoprotein expression vector. For MLV particles, cells were transfected with 500ng of MLV GagPol expression vector, 400ng of MLV-CMV-Cre, and 100ng of glycoprotein expression vector. For VSV particles, cells were transfected with 1 microgram of glycoprotein expression, and were infected 2 days post-transfection with >10^7^ infectious units/well of VSVΔG-GFP (Kerafast, (22)). Cells were rinsed with PBS one hour after infection and replace with complete media supplemented with 2 microliters of mouse hybridoma supernatant containing anti-VSV-G antibody I1 (Kerafast) to neutralize input virus. Neutralizing anti-body was excluded from samples pseudotyped with VSV-G. VSV pseudoparticles were collected 24 hours later.

Supernatant containing virus was frozen at -80°C for at least 4 h, thawed, and spun at 3,200 x *g* for 5 min, and the same volume of medium was added to target cells with 20 micrograms of hexadimethrine bromide per ml (H9268; Sigma). For assays with a fluorescent readout, infected cells were collected at about 2-3 days post-infection, fixed with 4% paraformaldehyde, washed with phosphate-buffered saline (PBS), and analyzed on an Accuri C6 flow cytometer. For infections with a Gluc readout, transductions were allowed to proceed for 2-3 days and 20 microliters of supernatant from each well was transferred to a black 96-well plate for measuring *Gaussia* luciferase activity with 50 microliters of 10 micromolar coelenterazine in 0.1 M Tris (pH 7.4) and 0.3 M sodium ascorbate (NanoLight Technology). Luminescence, representing infectivity, was measured from the supernatant using a PerkinElmer Enspire 2300 Multilabel Reader.

### Cell culture

The 293FT cell line was obtained from Invitrogen. All cells were maintained in Dulbecco’s modified Eagle’s medium (DMEM) supplemented with 10% fetal bovine serum, 2 mM L-glutamine, 1 mM sodium pyruvate, 10 mM nonessential amino acids, and 1% minimal essential medium (MEM) vitamins.

### Cell line generation

The Cre sensor cell line was generated by transfecting 293FT cells with 500 ng MLV GagPol expression vector, 400 ng of retroviral transfer vector MLV lox-mTomato-lox-GFP-Blast, and 100 ng of VSV-G expression vector. Viral media was used to transduce 293FT cells, and cells were selected with blasticidin (5 micrograms/ml) beginning 2 days post-transduction and maintained until control treated cells were all eliminated. A clonal isolate from these cells was selected that expressed mTomato but no GFP. The ACE2 cell lines were generated by transfecting 293FT cells with 500 ng MLV GagPol expression vector, 400 ng of retroviral transfer vector pQCXIP-ACE2 or pQCXIH-ACE2, and 100 ng of VSV-G expression vector. Viral media was used to transduce 293FT cells or the 293FT sensor cell line, and cells were selected with puromycin (1 microgram/ml) or hygromycin (200 micrograms/ml) beginning 2 days post-transduction and maintained until control treated cells were all eliminated. The TMPRSS2 cell line was generated by transfecting 293FT cells with 500 ng MLV GagPol expression vector, 400 ng of retroviral transfer vector pQCXIH-TMPRSS2, and 100 ng of VSV-G expression vector. Viral media was used to transduce 293FT, 293FT/ACE2, or 293/Cre-sensor/ACE2 cells were selected with hygromycin (200 micrograms/ml) beginning 2 days post-transduction and maintained until control treated cells were all eliminated.

### Neutralization assay

Deidentified patient plasma samples were obtained from the clinical laboratory MU healthcare. Plasma was heat-inactivated for 30 minutes at 58C. Two-fold serially diluted plasma was incubated with HIV-1/SARS-COV2 Spike pseudotypes for 1 hour at 37C. The mixture was subsequently incubated with 293/ACE2/TMPRSS2 cells, approximately 20,000 cells per well in a 96 well plate. Gluc measurements were taken 2-3 days post transduction.

## Discussion

Here we outline glycoprotein and target cell modifications that enhance pseudotyping efficiency with the SARS-COV2 Spike glycoprotein. The first and most important changes are the necessity to express the human ACE2 receptor, which has been defined previously by numerous investigators, and the enhancement gained from truncation of the Spike cytoplasmic tail. Truncation of the last 19 amino acids for SARS-COV1 was also shown to significantly enhance pseudotyping with that glycoprotein (9). In that study, the original reason for making the truncation was to eliminate potential ER retention sequences in the cytoplasmic tail. Despite enhancing pseudotyping capacity, the truncation did not enhance surface expression. It is likely that the enhancement in pseudotyping efficiency upon deletion of the cytoplasmic is at least in part due to eliminating steric interference of the cytoplasmic tail with the viral capsid. Such interference has been noted when pseudotyping other viral glycoproteins with large cytoplasmic tails, such as HIV-1 Env (23, 24).

The next modification that enhanced efficiency was the introduction of the human protease TMPRSS2. The importance of this protease for SARS-COV2 Spike glycoprotein entry has been noted previously, so this enhancement was not surprising (3, 6).

The final modification that enhanced SARS-COV2 Δ19 Spike pseudotyping was the introduction of the R682Q mutation, which is predicted to eliminate the furin cleavage site. This mutation was naturally selected for upon culturing SARS-COV2 in Vero cells, and viruses containing the mutation were reported to have larger plaque sizes (8). This observation is slightly at odds with a recent publication by Hoffmann et al. that found that the furin cleavage site was essential for infecting the lung cell line Calu-3, but not Vero cells. The Calu-3 cell line is reported to express TMPRSS2, but lack sufficient CatB/L to promote glycoprotein priming (3, 25); the Vero cells are reported to be TMPRSS2 negative, but CatB/L positive. Thus, the furin cleavage site appeared to only be required when the virus is dependent on TMPRSS2 for entry. A possible explanation is that furin cleavage enhances TMPRSS2 mediated entry, but suppresses CatB/L mediated entry. Consistent with this, the enhancement we observed from the R682Q mutation was less pronounced in cells that express TMPRSS2. It should be noted that mutations described in Hoffman et al. replaced the furin cleavage site from SARS-COV2 with the equivalent sequence from SARS-COV1, which effectively causes a 4 amino acid deletion. Thus, the loss of infectivity observed by Hoffmann et al. could in part have been the result secondary effects of the deletion that is independent of furin cleavage.

Finally, we demonstrate that HIV-1-Gluc particles pseudotyped with SARS-COV2 Δ19 Spike could be used for detecting neutralizing antibodies from COVID-19 patient plasma. Inclusion of the R682Q in Spike did not obviously change the neutralization results in this assay. Thus, this appears to be a suitable system for studying inhibitors of SARS-COV2 entry.

